## Supplementary Information for "Process-guidance improves predictive performance of neural networks for carbon turnover in ecosystems"

**1 Pipeline** **2**

**3 Architecture and hyperparameter search** **5**

**4 Technical details of the domain adaptation** **5**

**5 Technical details of the physics embedding** **8**

**6 Temporal predictions** **8**

**7 Bayesian Calibration of the process model** **9**

**References** **9**

### 1 Pipeline

#### 1.1 Structure

The pipeline structure is sketched in Fig. 1.

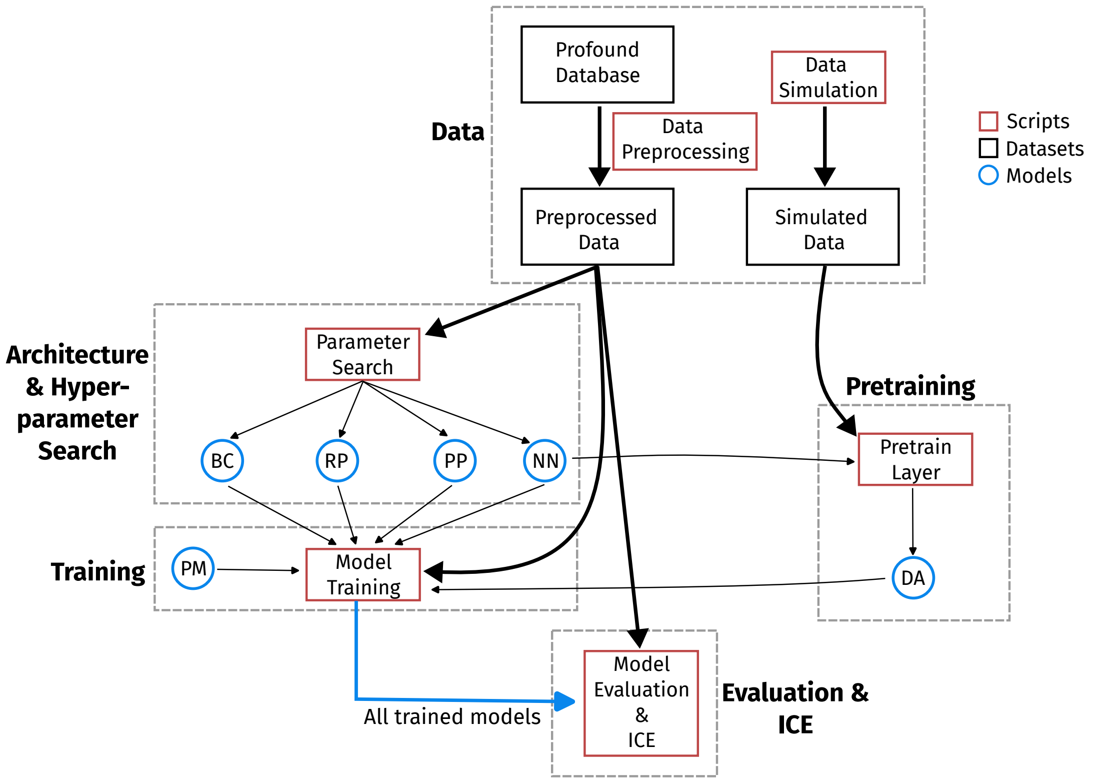

**Figure 1:** Sketch of the pipeline structure. The sequential order of the parts is from top to bottom. The structure is similar for the temporal and spatial experiment. PM = Process model, BC = Bias correction, RP= Residual physics, PP = Parallel physics, NN = Naive neural network, DA = Domain adaptation

#### 1.2 Download code

The code pipeline can be downloaded from GitHub `https://github.com/therealniklasmoser/physicsguidednn`.

**When using the pipeline code, cite the main article:**
Moser, N., Wesselkamp, M. and Dormann C. F. (2022). Process-guidance improves predictive performance of neural networks for carbon turnover in ecosystems

The pipeline consists of the folders src, data, code and r. The modified source code of PRELES is stored in the src folder. To compile the modified PRELES code to a PyTorch cpp-extension, that is callable from Python the source code needs to be compiled. We use the compiler version that is shown below.

```
niklas@hpc : $ gcc --version
gcc (GCC) 9.1.0
Copyright (C) 2019 Free Software Foundation, Inc.
This is free software; see the source for copying conditions.  There is NO
warranty; not even for MERCHANTABILITY or FITNESS FOR A PARTICULAR PURPOSE.
```

To convert the modified PRELES c-code to a proper PyTorch cpp-extension run the setup.py script in the src folder using the command:

```
niklas@hpc : $ python setup.py install
running install
running bdistegg
running egginfo
writing preles.egg-info/PKG-INFO
writing dependencylinks to preles.egg-info/dependencylinks.txt
writing top-level names to preles.egg-info/toplevel.txt
reading manifest file 'preles.egg-info/SOURCES.txt'
writing manifest file 'preles.egg-info/SOURCES.txt'
installing library code to build/bdist.linux-x8664/egg
running installlib
running buildext
building 'preles' extension
...
Processing dependencies for preles==0.0.0
Finished processing dependencies for preles==0.0.0
```

After running the setup script one can test if PRELES is callable in Python by the commands below. To call PRELES it is necessary to import torch first.

```
niklas@hpc : $ python
Python 3.7.10 — packaged by conda-forge — (default, Feb 19 2021, 16:07:37)
[GCC 9.3.0] on linux
Type "help", "copyright", "credits" or "license" for more information.

import torch
import preles
```

### 2 Data

#### 2.1 Source

We use the PROFOUND database. It provides information about the vegetation and climate at the forest stand scale in time series consisting of data of several years for forest sites in EuropeReyer et al. (2019). The PROFOUND database combines empirical data about forest stand structure, species composition, climatic conditions, energy and mineral fluxes. The data can be downloaded from the GitHub `https://github.com/COST-FP1304-PROFOUND/ProfoundData`.

#### 2.2 Preprocessing

For the analysis, the same input variables are used in the same units as in PRELES. The variables needed are namely the daily sums of photosynthetic active radiation (PAR), the mean air temperature (TAir), the vapour pressure deficit (VPD), the precipitation above canopy (Precip), the carbon dioxide concentration of air ($CO_2$), the fraction of photosynthetic active radiation (fAPAR) and the day of year (DOY), as depicted in Table 2. Additionally, information on the gross primary production (GPP) is required. It is used as the output variable in the models. The data required is in a daily resolution.

**Table 1:** Data used from the PROFOUND database.

| Abbreviation | Description | Unit |
|---|---|---|
| PAR | sum of photosynthetic active radiation | $\mathrm{mol\,m^{-2}\,d^{-1}}$ |
| TAir | mean air temperature | $^{\circ}\mathrm{C}$ |
| VPD | mean vapour pressure deficit | kPa |
| Precip | precipitation above canopy | mm |
| $CO_2$ | carbon dioxide concentration | ppm |
| fAPAR | fraction of photosynthetic active radiation | - |
| GPP | gross primary production | $\mathrm{g\,C\,m^{-2}\,d^{-1}}$ |

The PROFOUND database information on the irradiance, the global radiation $\phi_{\mathrm{e}}$ (in $\mathrm{J\,cm^{-2}\,d^{-1}}$) is converted into the quantum units describing the photon irradiance $\phi_{\mathrm{p}}$ (in $\mathrm{mol\,m^{-2}\,d^{-1}}$). Therefore, the relation of the wavelength and the energy of a photon is used as described in (Taiz & Zeiger 2015) with

$$\phi_{\mathrm{p}} = \frac{\phi_{\mathrm{e}}}{E N_{\mathrm{A}}} \quad , \tag{1}$$

and

$$E = \frac{hc}{\lambda} \quad , \tag{2}$$

where $h$ is Planck's constant ($6.63 \cdot 10^{-34}\mathrm{J\,s}$), $c$ is the speed of light ($2.99792458 \cdot 10^{8}\mathrm{m\,s^{-1}}$), $\lambda$ is the wavelength ($\approx 2.2 \cdot 10^{-7}\mathrm{m}$), $N_{\mathrm{A}}$ is Avogadro's constant ($6.602 \cdot 10^{23}\mathrm{mol^{-1}}$) and $\phi_{\mathrm{e}}$ is the global radiation (in $\mathrm{J\,s^{-1}\,m^{-2}}$).

fAPAR is derived from MODIS satellite data and has a resolution of eight days. The data gaps of the eight day resolution data are filled by averaging information of the data point before and after the gap. Second, to get the daily resolution, it is assumed that the eight day value is representative for all days within this period. Therefore, fAPAR is set constant over the eight days.

GPP is converted from $\mu\mathrm{mol\,CO_2\,m^{-2}\,s^{-1}}$ to $\mathrm{g\,C\,m^{-2}\,d^{-1}}$ using the molar mass of carbon ($\approx 12.011\mathrm{g\,mol^{-1}}$).

#### 2.3 Normalisation

Each variable is normalised around its mean $\sigma$ and its standard deviation $\sigma$ before using it as an input in the neural networks. Therefore, a $z$-score is calculated for each variable as

$$z = \frac{x - \mu}{\sigma} \quad . \tag{3}$$

Because of its cyclic character a sine and cosine function is used to normalise DOY ($d$). This ensures that the information of the last day of year $n$ and the first day of year $n + 1$ are closer together (compared to $365$ and $1$). DOY is transformed as

$$d_{\text{sin}} = \sin\left(d\frac{2\pi}{365}\right) \quad , \tag{4}$$

and

$$d_{\text{cos}} = \cos\left(d\frac{2\pi}{365}\right) \quad . \tag{5}$$

### 3 Architecture and hyperparameter search

We implemented a combined architecture and hyper-parameter search. The architecture search space consists of $300$ randomly sampled layersizes with a maximum depth of $4$ hidden layers. The hyper-parameter search space consists of $300$ randomly sampled hyper-parameter vectors. In the combined search each network of the $300$ architecture candidates is fitted using the $300$ candidate hyper-parameter vectors. Each hyper-parameter vector consists of learning rate, batch size. For the physics embedding and the physics regularisation the hyper-parameter vectors consist additionally of a regularisation factor. The best performing candidate network $k$ is chosen based on the minimum of the index $i_k$ with

$$i_k = \frac{\mathbf{E}\left[\mathcal{L}_{\text{val}}\right]^2 + \sqrt{\mathbf{E}\left[(\mathcal{L}_{\text{val}} - \mathbf{E}\left[\mathcal{L}_{\text{val}}\right])^2\right]}}{2} \quad , \tag{6}$$

where $\mathcal{L}_{\text{val}}$ denotes the validation losses for all cross-validation runs.

The best performing architecture and hyper-parameters are shown for each model in Tab. 3 for the full data experiment and in Tab. 3 for the sparse data experiment.

The performance of the candidate architectures and hyper-parameters are shown in Fig. 2 for the temporal experiment.

### 4 Technical details of the domain adaptation

The yearly equivalents of the neural network training data were the basis for the climate simulations. Except for $CO_2$, which was fixed at a value of 380 ppm, the five climatic variables $y = \{$T, $\phi$, D, P, $f_{\text{aPPFD}}\}$ were separately modeled with a generalized additive model (GAM). For the on-site simulations, each $y_k \in y$ was described as a function of the day

**Table 2:** Full data architecture and hyper-parameters for each model and the temporal experiment

| Model | Architecture | LR | BS | $\lambda$ | $i$ |
|---|---|---|---|---|---|
| NN | $[2, 64, 128]$ | 0.0143 | 16 | - | 0.306126 |
| BC | $[8, 64]$ | 0.0531 | 16 | - | 2.928012 |
| PP | $[256, 128]$ | 0.002 | 16 | - | 1.769507 |
| RP | $[256, 32, 8]$ | 0.0265 | 4 | 0.0101 | 0.584885 |

**Table 3:** Sparse data architecture and hyper-parameters for each model and the temporal experiment

| Model | Architecture | LR | BS | $\lambda$ | $i$ |
|---|---|---|---|---|---|
| NN | $[2, 8, 64]$ | 0.0082 | 2 | - | 0.943277 |
| BC | $[16, 256]$ | 0.0122 | 4 | - | 1.371335 |
| PP | $[2]$ | 0.0735 | 64 | - | 3.362334 |
| RP | $[2, 256, 8, 256]$ | 0.0184 | 16 | 0.0654 | 1.354255 |

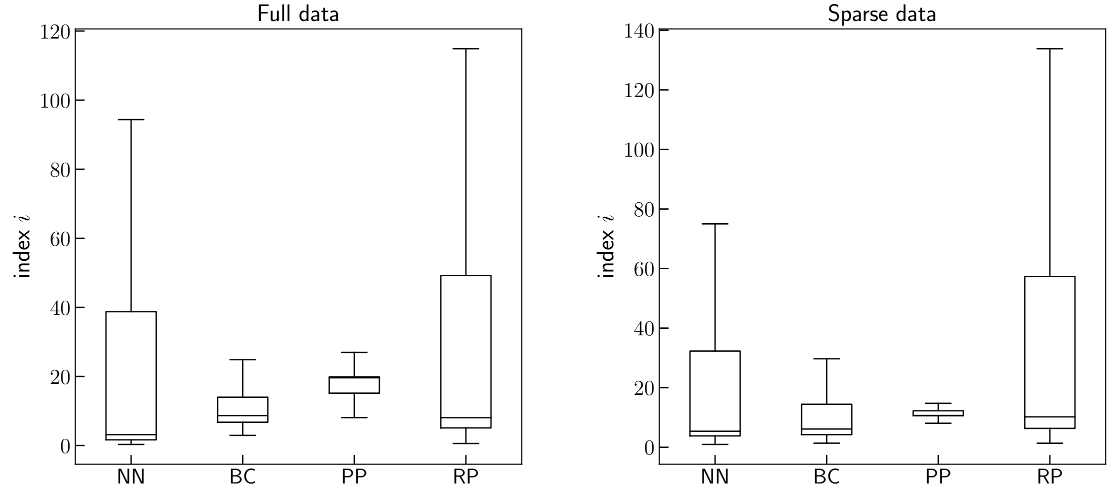

**Figure 2:** Index values for all candidate models in the architecutre and hyper-parameter search. PM = Process model, BC = Bias correction, RP= Residual physics, PP = Parallel physics, NN = Naive neural network, DA = Domain adaptation

**Table 4:** $R^2$ statistics of fitted GAMs for synthetic data generation under sparse data assumption.

| Variable | On-site $R^2$ | Multi-site $R^2$ |
|---|---|---|
| $T$ | 0.9836 | 0.7427 |
| $R$ | 0.9791 | 0.3995 |
| $D$ | 0.9582 | 0.7145 |
| $\phi$ | 0.9928 | 0.7888 |
| $f_{\text{aPPFD}}$ | 0.9771 | 0.6761 |

of the year (DOY) in interaction with the year $R$. The GAM was fitted with a cyclic cubic smooth function $f$ imposed onto DOY. Allowing for the interaction with $R$, the smooth varies with the year.

$$y_k = \beta_0 + \sum_{i=1}^{4} f_i(\text{DOY}) \cdot R_i + \epsilon \tag{7}$$

For the multi-site simulations, each $y_k$ was described as a function of the day of the year (DOY) in interaction with the site $S$.

$$y_k = \beta_0 + \sum_{i=1}^{5} f_i(\text{DOY}) \cdot S_i + \epsilon \tag{8}$$

The resulting models could be used to simulate estimated time series of the variables of any length and for any time point of years and sites under consideration. To introduce variation to the simulation, noise $\nu$ was added to each $\hat{y}_k$, sampled from a multivariate normal distribution $\nu \sim \mathcal{N}_{k=5}(\mu, \Sigma)$. Here, $\mu$ denotes the $k$-dimensional mean vector and $\Sigma$ is the $k \times k$-dimensional covariance matrix. Means $\mu$ were set to zeros while the covariance matrix $\Sigma$ was specified explicitly as the covariance matrix of the $k$-dimensional residual matrix.

With both, a global and local sensitivity analyses, the five most sensititive stand specific PRELES parameters were considered. These are the potential light use efficiency ($\beta$), the threshold for the state of acclimation ($X0$), the light modifier parameter for saturation with irradiance ($\gamma$), and the transpiration and evaporation parameters ($\alpha$ and $\chi$). Prior knowledge about their marginal distributions is available as uniform prior distribution parameters, used for the Bayesian calibration of PRELES (Minunno et al. 2016). In order to provide the neural network with the same information as PRELES, the prior distributions were strongly narrowed down, ressembling the default calibration. In a Latin hypercube design the five parameters were sampled from their neat uniform joint probability distribution. The other parameters remained fix at their default values.

### 5 Technical details of the physics embedding

We convert the PRELES c-code into a callable Python library. Therefore, we change the elementary operations in the c-code to their PyTorch equivalent, e.g. `sin(x)`→`torch.sin(x)`. This is a necessary condition to use PRELES as a forward pass of the neural network. In the PyTorch framework all calculations on the input tensor are stored in a computational graph during forward propagation. During backpropagation the computational graph of the tensor is used to numerically calculate intermediate gradients for all elementary operations on the tensor.

### 6 Temporal predictions

The temporal predictions with the full data set are shown in Fig. 3.

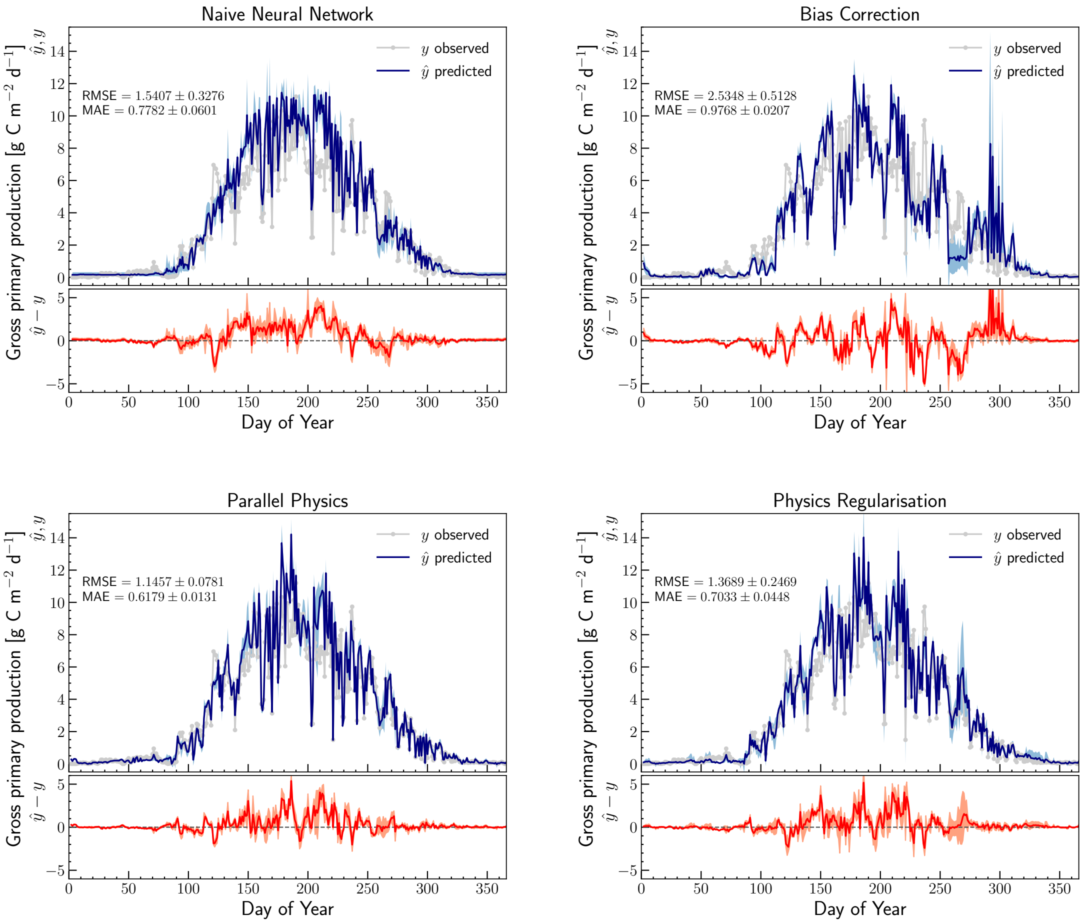

**Figure 3:** Temporal predictions with the full data set.

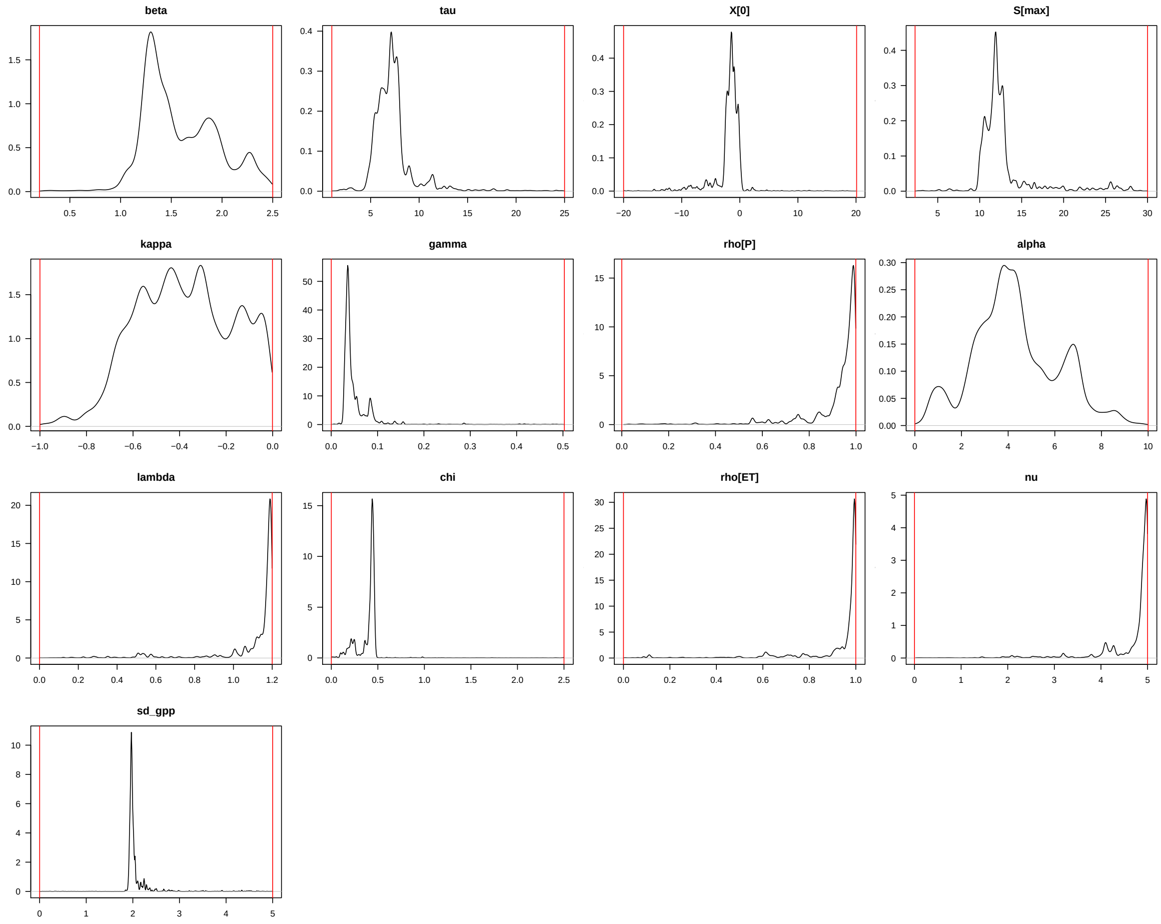

**Figure 4:** Posterior distributions of the re-calibrated PRELES parameters.

### 7 Bayesian Calibration of the process model

PRELES was re-calibrated using the Bayesian Tools package (see main document). A Markov Chain Monte Carlo simulation using the DREAMzs sampler was used. We ran three chains at 50000 iterations each. The results are shown in Fig. 4.
